# Multiscale three-dimensional ultrastructural mapping of intestinal tissues and organoids

**DOI:** 10.64898/2026.06.30.734790

**Authors:** Thijs Makaske, Vincent Hellebrekers, Albert K. Serweta, Dimitrios Laskaris, Saskia J.E. Suijkerbuijk, Sabine A. Fuchs, Kerstin Schneeberger-Verjaal, Jacco van Rheenen, Ihor Smal, Lukas C. Kapitein

## Abstract

Understanding cell biology in native environments requires imaging of subcellular organization in three dimensions. In the intestinal epithelium, multiple cell types organize along the crypt-villus axis, where cell-cell interfaces and subcellular architecture control cell differentiation, tissue organization and epithelial function. Resolving these features volumetrically remains challenging: light microscopy offers molecular specificity but has limited resolution, whereas electron microscopy provides ultrastructural detail but is poorly suited to volumetric acquisition combined with specific protein labeling. Here, we show that expansion microscopy enables the multiscale volumetric study of epithelial ultrastructure in tissue sections and organoid models. Using an optimized workflow, we resolve epithelial tissue architecture, cell types and subcellular features within volumes across scales. Application to a microvillus inclusion disease (MVID) organoid model revealed disease-associated ultrastructural phenotypes that were only observed using electron microscopy. Our results establish expansion microscopy as key technology for studying three-dimensional cell biology within intestinal tissue and tissue mimics.

## INTRODUCTION

Epithelia are polarized tissues whose three-dimensional organization is fundamental to the structure and function of many organs. Cell-cell interactions, cellular morphology, and apico-basal polarity are tightly coordinated to support tissue architecture and physiological function.^1,2^ The intestinal epithelium is a striking example of this organization, containing a continuous differentiation axis from intestinal stem cells (ISCs) at the base of the crypt to differentiated cells toward the villus tip.^3^ Throughout this crypt-villus axis, epithelial cells establish highly polarized apical and basolateral membrane domains, organized brush borders, junctional complexes, and extensive lateral membrane contacts. These are essential for nutrient absorption, barrier function, and stem cell niche organization.^4^

Intestinal organoids have emerged as powerful stem cell-derived model systems that recapitulate many key features of epithelial tissues, including multicellular organization, differentiated cell types, and apico-basal polarity, thereby enabling cell biological analysis of differentiated epithelial cell types in vitro.^5^ Validation of organoids as physiologically relevant tissue models is commonly based on omics approaches such as single-cell RNA sequencing,^6^ while ultrastructural validation remains largely restricted to sparse two-dimensional electron microscopy images.^5^ However, epithelia are inherently three-dimensional structures, making high-resolution volumetric imaging approaches critical to validate tissue mimics and to explore epithelial cell biology.

Despite the need for volumetric visualization, imaging the intestine in three dimensions across scales remains challenging. Conventional fluorescence microscopy enables imaging specific molecules within intact tissue and organoids but is limited by optical scattering and relatively poor axial resolution. In contrast, electron microscopy (EM) provides substantially higher spatial resolution and remains the gold standard for ultrastructural analysis. However, EM typically requires complex sample preparation workflows, offers limited compatibility with multiplexed molecular labeling, and remains difficult and time-consuming to scale toward large tissue volumes.^7^ As a result, tissue-scale organization and subcellular ultrastructure are frequently analyzed separately using different imaging modalities.^8^

Expansion microscopy (ExM) embeds biological samples within hydrogels that can swell to physically enlarge the sample and enable fluorescence microscopy to resolve features beyond the conventional diffraction limit.^9,10^ Ultrastructure expansion microscopy (U-ExM) further improves preservation of subcellular organization and has enabled visualization of cytoskeletal and organellar ultrastructure in a variety of biological systems.^11–13^ Combined with dense NHS-ester total protein labeling^14,15^, U-ExM offers the possibility to bridge volumetric fluorescence imaging with ultrastructural analysis while remaining compatible with molecularly specific labeling strategies. These properties make U-ExM particularly attractive for studying intestinal epithelial systems, where tissue organization, epithelial topology, and subcellular architecture should be interpreted simultaneously within three-dimensional volumes.

Here, we establish an optimized expansion microscopy workflow for epithelial tissue and organoids that enables multiscale imaging from tissue architecture to subcellular ultrastructure within intact epithelial systems. By optimizing fixation, gel monomer infiltration, and homogenization, we achieve isotropic expansion while preserving sample integrity across large tissue volumes. Combined with total protein labeling, targeted antibody staining, and volumetric segmentation, this framework enables molecularly annotated three-dimensional reconstruction of intact intestinal crypts at single-cell resolution. Using this approach, we visualize epithelial topology, reconstruct complete crypt organization, as well as disease-associated ultrastructural phenotypes in a microvillus inclusion disease (MVID) organoid model. Together, our results establish the U-ExM workflow as a platform for quantitative multiscale imaging of intact epithelial systems.

## RESULTS

### Ultrastructure expansion microscopy enables multiscale imaging of mouse small intestine

To resolve intestinal architecture across spatial scales, we established an ultrastructure expansion microscopy workflow for mouse small intestine tissue sections (**Fig. 1A**). To improve preservation of epithelial ultrastructure, tissue sections were fixed in 4% paraformaldehyde (PFA) in PHEM buffer, a fixation strategy commonly used for preserving cytoskeletal architecture and ultrastructure.^16,17^ In addition, inactive and activated gelation solutions were incubated for extended periods relative to the original U-ExM protocol to improve monomer penetration throughout dense tissue volumes. To improve tissue homogenization and isotropic expansion, samples were denatured in an autoclave at 126°C for 2 hours (**Sup. Fig. 1**).^18^ This protocol enabled robust expansion of dense epithelial tissue sections.

**Figure 1.**
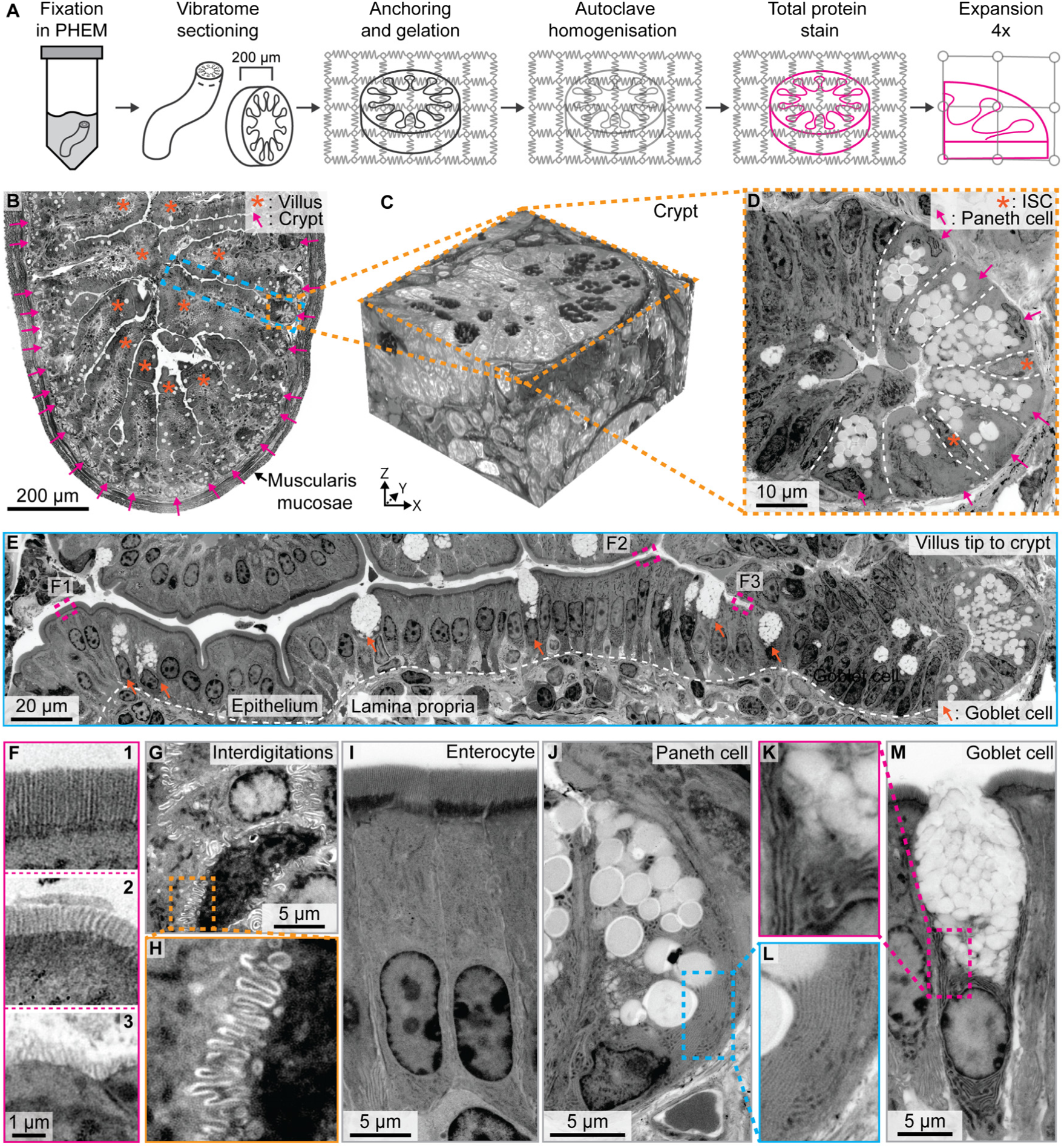
Ultrastructure expansion microscopy of mouse small intestine tissue. **A)** A schematic overview of the optimized expansion workflow for mouse small intestine tissue slices. **B)** Tissue slice of a mouse small intestine showing multiple crypts (arrow) and villi (asterisk) embedded within the lamina propria and surrounded by the muscularis mucosae. **C)** Three-dimensional rendering of an intestinal crypt. **D)** Single crypt highlighting its stem cell niche, including Paneth cells (arrow) and adjacent intestinal stem cells (asterisk) at the crypt base. **E)** Continuous view from villus tip to crypt showing the intestinal lamina propria and epithelium with goblet cells (arrow) in between enterocytes. **F)** Gradual changes in microvillar organization along the villus-crypt axis, highlighting spatial variations in brush border maturation. F1 at the villus tip to F3 at the villus base. **G)** Lateral interdigitations between neighboring enterocytes, resolved at subcellular scale. **H)** Higher-magnification view of (**G**), revealing fine membrane interlocking structures. **I, J, M**) Major epithelial cell types identified within the organoid epithelium: enterocyte (**I**), Paneth cell (**J**), and goblet cell (**M**). **K)** High-resolution view of goblet cells (from **M**) showing protein dense compartments. **L)** High-resolution view of Paneth cells (from **J**) highlighting protein dense compartments. Scale bars are corrected for expansion factor. All images show the NHS-ester Atto-643 total protein stain.

The workflow preserved tissue-scale organization after expansion, allowing imaging of multiple continuous crypt-villus units in a single stitched overview (**Fig. 1B and Sup. Video 1**). Individual crypts could be reconstructed in three dimensions, maintaining continuous epithelial topology across large volumes (**Fig. 1C**). At higher resolution, the protocol enabled clear visualization of the crypt niche, including Paneth cells and the adjacent, narrow ISCs (**Fig. 1D**). Moreover, the optimized workflow enabled continuous imaging across intact crypt-villus units while preserving epithelial continuity and tissue architecture (**Fig. 1E**).

At subcellular resolution, brush border organization varied along the crypt-villus, with more densely packed and ordered microvilli toward the villus and less organized structures near the crypt (**Fig. 1F**). At the lateral membrane, interdigitations between neighboring enterocytes were resolved (**Fig. 1G**), forming complex interlocking structures visible at higher magnification (**Fig. 1H**). Major epithelial cell types were identifiable based on morphology, including enterocytes, goblet cells with protein dense organelles, and Paneth cells with prominent granules **(Fig. 1I-M** and **Sup. Fig. 2)**. Together, these data show that our optimized ExM workflow enables multiscale imaging of intact mouse small intestine tissue, linking tissue architecture to subcellular ultrastructure within a single framework.

**Figure 2.**
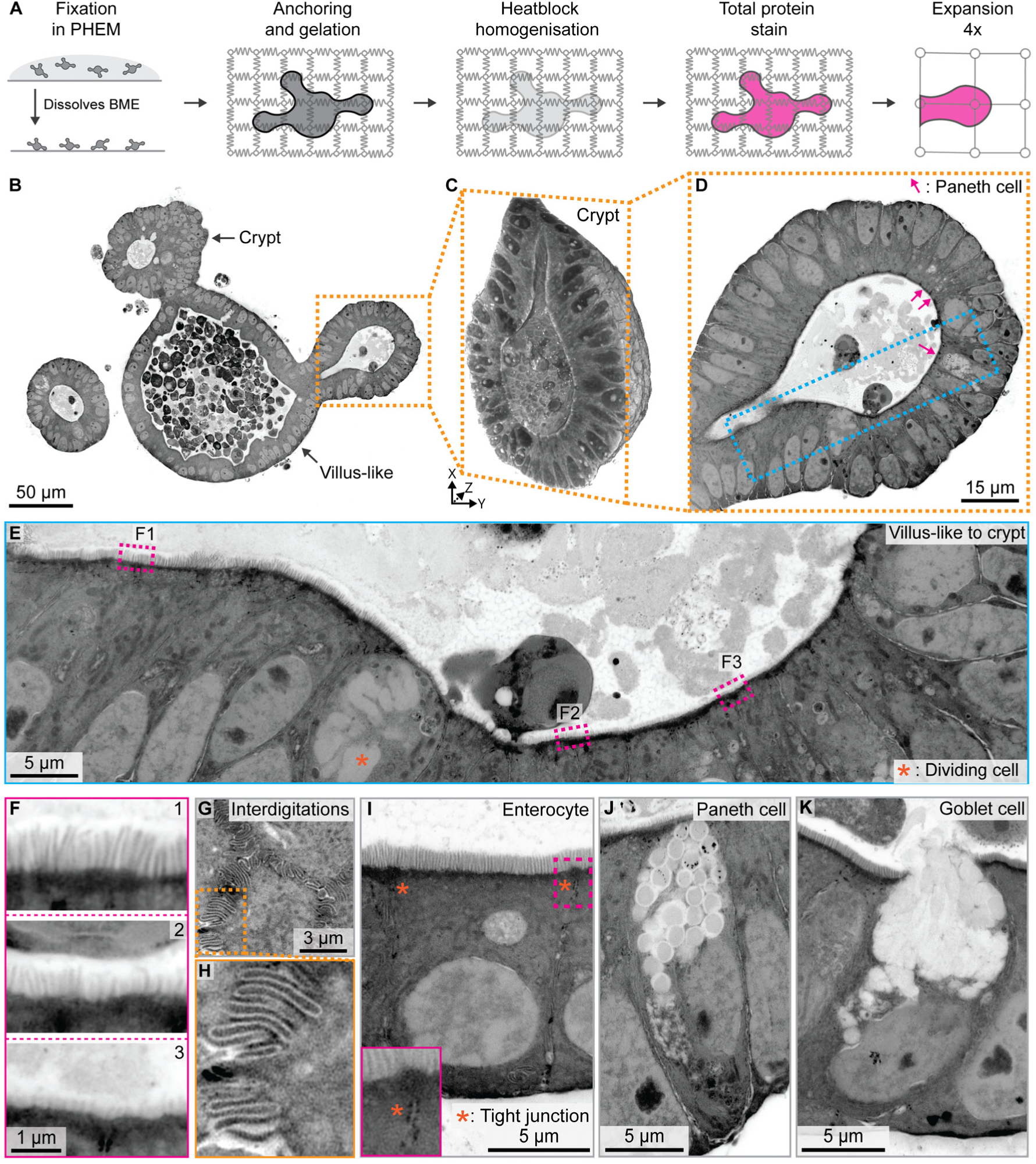
Ultrastructure expansion microscopy of mouse small intestine organoids. **A)** A schematic overview of the optimized expansion workflow for mSI organoids. **B)** Optical section through an intact mSI organoid revealing multiple crypt domains and a continuous epithelial architecture at subcellular resolution. **C)** Three-dimensional rendering of a single crypt from (**B**), preserving full epithelial morphology. **D)** Single crypt highlighting Paneth cells (arrow) localized to the crypt base. **E)** Continuous view of brush border organization along the crypt axis, including a dividing cell (asterisk). **F)** Gradual changes in microvillar organization along the villus-crypt axis, highlighting spatial variations in brush border maturation. F1 at near the villus-like region to F3 at the crypt base. **G)** Lateral interdigitations between neighboring enterocytes. **H)** Higher-magnification view of (**G**), revealing fine membrane interlocking structures. **I, J, K**) Major epithelial cell types identified within the organoid epithelium: enterocyte (**I**) with a zoom-in highlighting tight junctions, Paneth cell (**J**), and goblet cell (**K**). Scale bars are corrected for expansion factor. All images show the NHS-ester Atto-643 total protein stain.

### Ultrastructure expansion microscopy enables multiscale imaging of mouse small intestine organoids

Having established that the optimized ExM workflow preserves tissue architecture and ultrastructure in mouse small intestine tissue, we next asked whether this approach could be extended to organoid systems, which recapitulate key aspects of intestinal epithelial organization in vitro, including multicellular architecture, differentiated cell types and apico-basal polarity. Because organoid ultrastructural characterization has so far mostly remained limited to two-dimensional electron microscopy images, we reasoned that volumetric ultrastructural imaging could provide an additional framework for evaluating epithelial organization across native tissue and organoid systems.

We adapted our workflow for mouse small intestine (mSI) organoids by replacing autoclave-based homogenization with heat-block incubation, which was sufficient to achieve complete tissue homogenization while preserving epithelial organization and enabling uniform expansion and total protein labeling (**Fig.2A**). Similar to the tissue sections, fixation was performed using PHEM buffer, which improved preservation of epithelial integrity and cellular ultrastructure compared to PBS-based fixation conditions that were initially used (**Sup. Fig. 3**). Expanded organoids retained their overall architecture, revealing multiple crypt-like domains within a continuous epithelial structure at subcellular resolution (**Fig. 2B**). Three-dimensional reconstruction of individual crypts enabled direct visualization of crypt organization in intact organoids and revealed that epithelial topology was preserved over large volumes (**Fig. 2C, D**).

**Figure 3.**
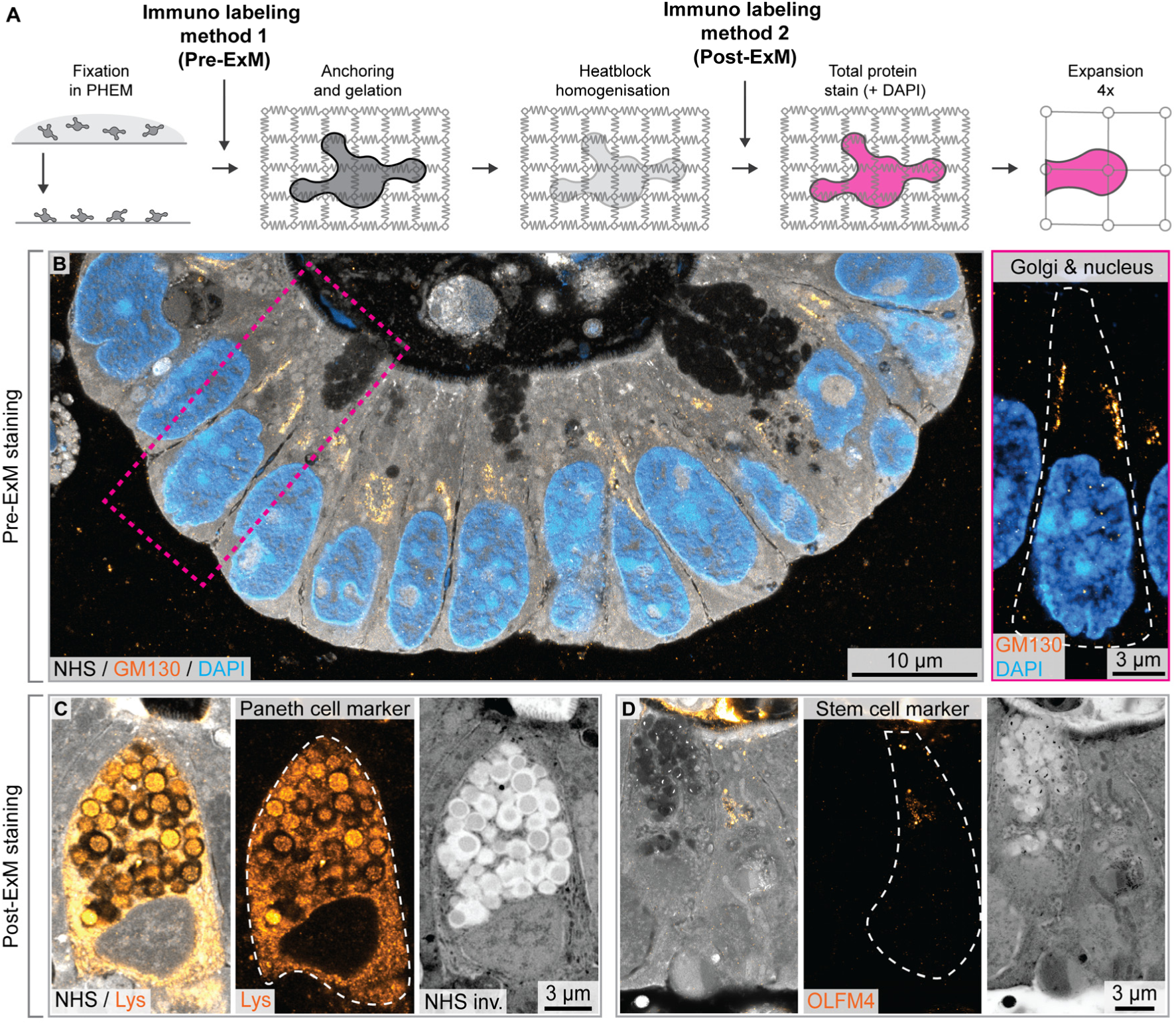
Pre- and post-ExM targeted antibody labeling. **A)** Schematic overview indicating the position of pre- and post-expansion antibody labeling within the ExM workflow. **B)** Pre-expansion antibody labeling against GM130, a Golgi marker, combined with NHS-ester Atto-643 total protein staining and DAPI. Right panel shows a higher-magnification crop displaying multiple Golgi’s near the lateral membranes. **C)** Post-expansion antibody labeling against lysozyme, identifying Paneth cells within the organoid epithelium, together with NHS-ester Atto-643 total protein staining. **D)** Post-expansion antibody labeling against OLFM4, identifying stem cells at the crypt base, together with NHS-ester Atto-643 total protein staining. Scale bars are corrected for expansion factor.

Continuous imaging along the crypt axis revealed progressive changes in epithelial ultrastructure (**Fig. 2E**). Microvillar organization varied along this axis, with more densely packed and ordered brush border structures observed toward villus-like regions and less organized microvilli near the crypt base (**Fig. 2F**), consistent with spatial differences in epithelial maturation and similar to what is observed in the native tissue (**Fig. 1F**). Dividing cells could also be identified within the crypt region (**Fig. 2E**). At the lateral membrane, expansion microscopy resolved interdigitating contacts between neighboring enterocytes (**Fig. 2G, H**). Major epithelial cell types were readily identifiable based on morphology and intracellular organization, including enterocytes with defined tight junctions, goblet cells, and Paneth cells with prominent granules (**Fig. 2I-K**). Together, these results reveal that our ExM workflow enables multiscale volumetric imaging of intestinal organoids while preserving key architectural and ultrastructural features also observed in native intestinal epithelium, thereby enabling direct comparison of epithelial organization across native and in vitro systems.

### Compatibility with pre- and post-ExM targeted antibody labeling

To enable molecular annotation of ultrastructural datasets, we integrated both pre-and post-expansion antibody labeling into the U-ExM workflow (**Fig. 3A**). Pre-expansion labeling enables staining of epitopes prior to chemical and structural modification during expansion processing, whereas post-expansion labeling reduces effective linkage error between fluorophore and target epitope.

Pre-expansion labeling against the Golgi marker GM130 was preserved following expansion and could be visualized together with NHS-ester total protein staining and DAPI (**Fig. 3B**). An enlarged view revealed cells containing multiple Golgi positioned near the lateral membranes and adjacent to a Paneth cell, consistent with recent observations obtained using serial block-face scanning electron microscopy showing that ISCs spatially orient and partition their Golgi apparatus relative to niche contacts.^19^

We next tested post-expansion antibody labeling using cell type–specific markers within intact organoids. Post-expansion staining against Lysozyme identified Paneth cells within the crypt epithelium while maintaining overall structural context through NHS-ester labeling (**Fig. 3C**). Similarly, post-expansion labeling against OLFM4 resolved ISCs localized at the crypt base (**Fig. 3D**). Thus, our U-ExM workflow is compatible with targeted antibody labeling both before and after expansion, enabling molecular annotation of ultrastructural datasets within intact intestinal organoids.

### ExM enables nanoscale segmentation and quantification of all cells in a crypt

Having established that our ExM workflow enables multiscale imaging of intact intestinal tissue and organoids, we next asked whether these datasets could support volumetric segmentation of the entire crypt epithelium into individual cell bodies for morphological and quantitative analyses. To achieve this, we combined NHS-ester total protein labeling with post-expansion antibody staining against membrane-localized CAAX-tdTomato and implemented a custom human-in-the-loop segmentation workflow to reconstruct all cells within an entire intestinal organoid crypt (**Fig. 4A and Sup. Video 2**). Three-dimensional rendering of the segmented dataset shows individual nuclei and complete cell bodies throughout the crypt epithelium (**Fig. 4B**). Overlay of the full cell body segmentation with the NHS channel demonstrated continuous reconstruction of the epithelial layer at single-cell resolution across the entire crypt volume without segmentation gaps (**Fig. 4C and Sup. Video 3**).

**Figure 4.**
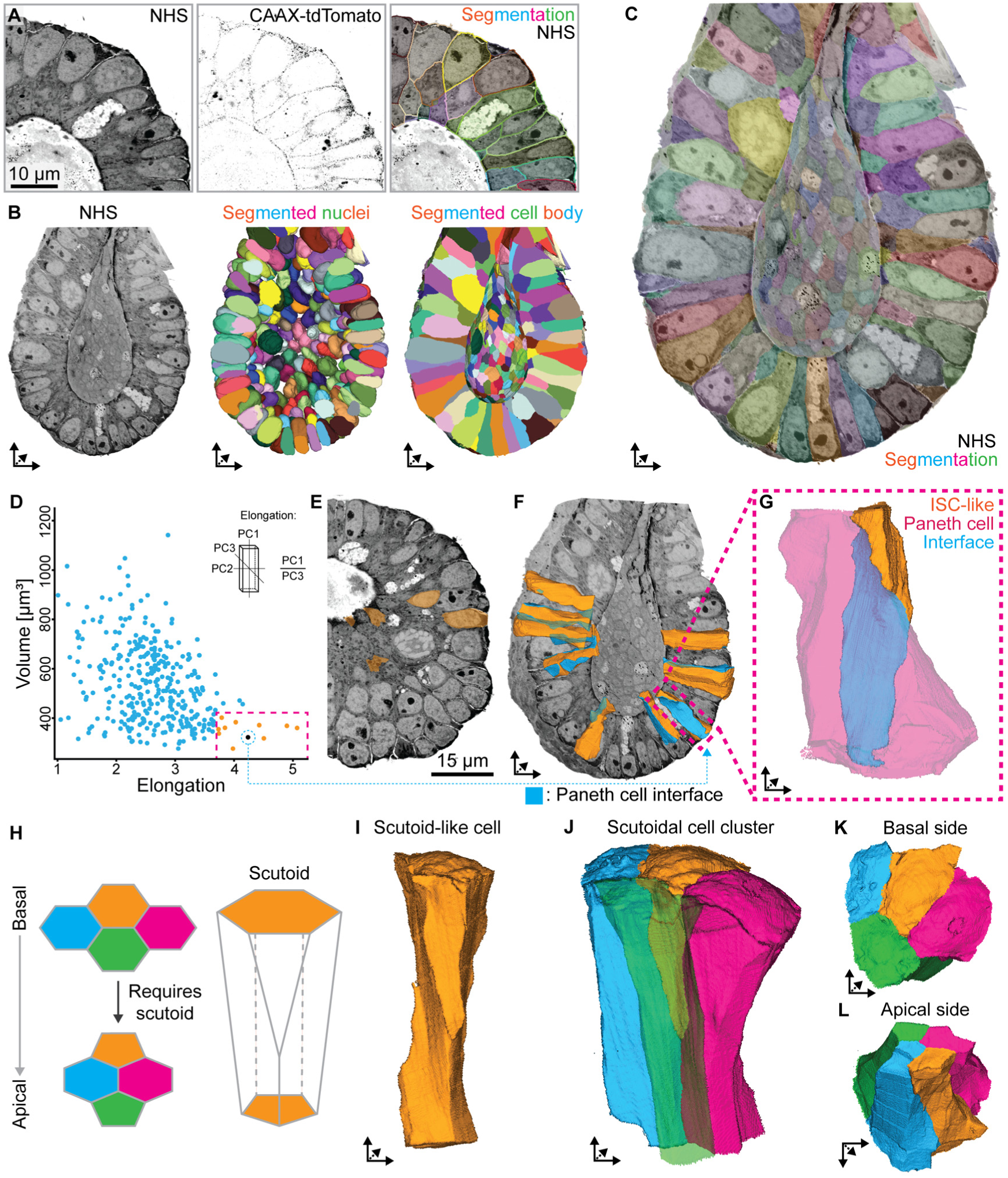
Volumetric segmentation of an entire intestinal organoid crypt. **A)** Two-dimensional view of the raw NHS-ester Atto-643 total protein stain, raw post-ExM antibody stain against CAAX-RFP and resulting human-in-the-loop segmentation as overlaid on the total protein channel. **B)** Three-dimensional rendering of the crypt showing the raw total protein signal, segmented nuclei, and segmented cell bodies. **C)** Volumetric rendering of the NHS channel overlaid with the full cell body segmentation, illustrating complete epithelial reconstruction at single-cell resolution. **D)** Quantification of cell volume versus elongation identifies a subset of small, highly elongated cells (magenta highlighted gate and orange individual cells). **E)** Slice through of the crypt showing apical cross-sections of elongated cells (orange), which are difficult to resolve in two-dimensional views, overlaid on the total protein channel. **F)** Three-dimensional rendering of elongated cells (orange) highlighting their extensive interfaces with Paneth cells (cyan). **G)** Higher-magnification view of an elongated cell wrapping around a Paneth cell, showing a large contact interface (cyan). **H)** Schematic representation of a scutoid, a geometrical cell shape enabling apico-basal neighbor exchange. **I)** Example of one of the scutoid-like cells identified within the segmented crypt. **J)** The scutoid-like cell shown in (**I**) within its local cellular neighborhood. **(K–L)** Basal (**K**) and apical (**L**) membrane surfaces of the cell cluster in (**J**), illustrating neighbor rearrangements consistent with scutoid topology. Voxel size is ∼80 nm corrected for expansion factor.

Quantification of cell volume and elongation identified a distinct subset of small, highly elongated ISC-like cells within the crypt epithelium (**Fig. 4D and Sup. Video 4**). In two-dimensional sections these cells were difficult to identify due to their limited cross-sectional area in individual sections (**Fig. 4E**), highlighting the need for three-dimensional data acquisition in these models. Moreover, three-dimensional reconstruction revealed that these elongated cells formed extensive interfaces with neighboring Paneth cells (**Fig. 4F**). Higher magnification demonstrated that some elongated cells partially wrapped around Paneth cells, generating large contact surfaces between the two cell types (**Fig. 4G**).

The morphology of some cells resembled scutoids, a recently identified geometrical shape associated with apico-basal neighbor exchange in curved epithelia (**Fig. 4H**).^20^ Consistent with this topology, segmented cells within the crypt displayed distinct apical and basal neighbor relationships (**Fig. 4I-L and Sup. Video 5**). Such apico-basal neighbor exchange has been proposed to expand the range of contact-dependent signaling interactions within curved epithelia by increasing the number of neighbors contacted across the apico-basal axis.^21^ Indeed, comparison of apical and basal membrane surfaces revealed neighbor rearrangements across the apico-basal axis. These geometrical features, including sharp interface transitions (**Fig. 4I**), complex three-dimensional cell packing (**Fig. 4J**) and distinct basal and apical contact pattern (**Fig. 4K&L**), were accurately captured by the cell segmentation. Together, these results show that ExM datasets enable nanoscale volumetric segmentation and quantitative reconstruction of intact intestinal crypts, enabling analysis of epithelial topology and cell morphology within complex three-dimensional tissue organization.

### U-ExM resolves disease-associated ultrastructural defects in MVID organoids

To determine whether the U-ExM workflow could visualize disease-associated epithelial ultrastructure, we applied the method to a microvillus inclusion disease (MVID) organoid model based on an inducible knockout of Myosin VB (*MYO5B*) (**Fig. 5A**). Loss of MYO5B is associated with disruption of apical trafficking, formation of microvillus inclusions, and reduced brush border organization. Detection of these hallmark ultrastructural phenotypes have so far required electron microscopy.^22–24^

**Figure 5.**
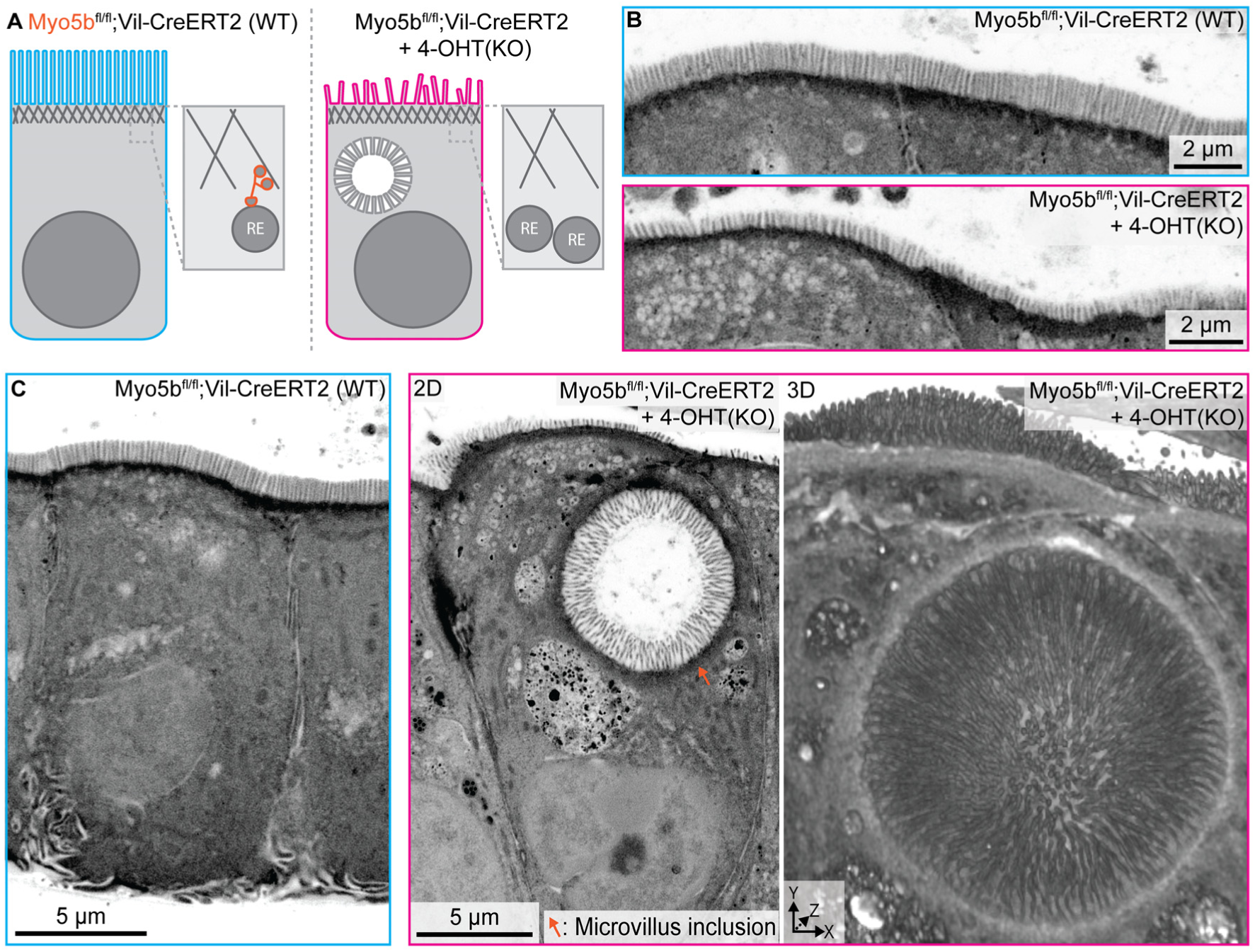
Volumetric ultrastructure of MVID organoids. **A)** Schematic of the inducible MVID model based on inducible *Myo5b* knockout, leading to the formation of microvillus inclusions and reduced microvilli. **B)** Apical brush border in non-induced cells (top) compared to 4-OHT induced *Myo5b* knockout cells (bottom), showing reduced apical microvilli and accumulation of subapical vesicles. **C)** Representative non-induced cell (left) and induced cell (middle) displaying a microvillus inclusion, subapical vesicles and large (auto)lysosomes, with a corresponding three-dimensional rendering of the induced cell (right). Scale bars are corrected for expansion factor. All images show the NHS-ester Atto-643 total protein stain.

In non-induced organoids, enterocytes displayed a continuous apical brush border with densely organized microvilli (**Fig. 5B, top**). In contrast, 4-OHT-induced *Myo5b* knockout organoids exhibited reduced apical microvilli together with accumulation of subapical vesicular structures and large (auto)lysosomes that are characteristic for the disease (**Fig. 5B, bottom**), consistent with previously described MVID phenotypes.^4,23^ Higher magnification imaging identified intracellular microvillus inclusions within induced cells (**Fig. 5C and Sup. Video 6**). Three-dimensional rendering of affected cells revealed the volumetric organization of these inclusions within the epithelial context of the organoid (**Fig. 5C, right**). Together, these results demonstrate that U-ExM enables volumetric visualization of disease-associated ultrastructural defects within intact intestinal organoids, extending the applicability of the workflow to pathological epithelial states.

## DISCUSSION

Resolving cell biology within tissues requires imaging approaches that bridge subcellular organization with tissue-scale architecture. This challenge is particularly acute in epithelial systems, where cellular morphology, apico-basal polarity, and cell-cell interactions are tightly integrated across three-dimensional space to support tissue function. Consequently, key structural features including membrane topology, lateral cell-cell interfaces, and polarized subcellular organization can be incompletely represented or misinterpreted in two-dimensional sections, especially within densely packed epithelia. The intestinal epithelium exemplifies this complexity. It contains a continuous crypt-villus axis in which ISCs transition toward differentiated enterocytes and goblet cells, while simultaneously establishing highly specialized membrane domains, junctional architectures, and brush border organization. Capturing these coordinated structural features therefore requires volumetric imaging approaches capable of resolving epithelial organization across spatial scales within intact three-dimensional tissue contexts.

Here, we established an improved expansion microscopy workflow for mouse small intestine tissue and organoids that enables multiscale imaging, molecular annotation, and quantitative three-dimensional reconstruction of intact intestinal epithelium using fluorescence microscopy. By optimizing fixation buffer, gel monomer infiltration, and homogenization, we achieved fourfold expansion while preserving tissue architecture and subcellular organization. Combined with NHS-ester total protein labeling, this revealed continuous visualization of the intestine tissue volumes from crypt-villus organization down to epithelial ultrastructure. This allowed a direct volumetric comparison between native tissue and organoid model systems, providing an additional layer of structural validation beyond transcriptional profiling approaches alone. ExM is particularly well suited for volumetric epithelial imaging because sample expansion simultaneously improves axial resolution, partially clears dense tissue, preserves ultrastructural organization, and remains compatible with molecularly specific labeling.

The compatibility of the workflow with pre- and post-expansion antibody labeling substantially extends its applicability. Pre-expansion labeling enabled detection of epitopes prior to chemical and structural modification during expansion processing, whereas post-expansion labeling reduced effective linkage error within expanded samples. Although many antibodies lose compatibility following post-expansion processing, physical decrowding of epitopes can improve labeling density in expanded samples.^25^ In our workflow, tertiary staining was incorporated into post-expansion labeling to further enhance signal intensity within expanded gels. Together, these approaches enabled molecular annotation of epithelial ultrastructure across multiple spatial scales.

By integrating ExM with a custom human-in-the-loop segmentation workflow, we reconstructed the complete crypt epithelium at single-cell resolution, enabling quantitative analysis of epithelial morphology and topology within intact tissue context. As demonstrated by the elongated crypt cells identified in our datasets, cellular morphology and neighbor relationships can be substantially underrepresented in two-dimensional sections due to limited apical cross-sectional area. Volumetric reconstruction further revealed extensive Paneth cell interfaces and scutoid-like geometries associated with apico-basal neighbor exchange, illustrating how three-dimensional imaging can reveal complex epithelial organization that is difficult to accurately interpret in conventional two-dimensional sections.

We further demonstrate that the workflow can resolve disease-associated ultrastructural phenotypes in an inducible *Myo5b* knockout organoid model of microvillus inclusion disease (MVID). In addition to reduced brush border organization, ExM visualized intracellular microvillus inclusions, accumulation of subapical vesicular structures and large (auto)lysosomes within intact epithelial context, illustrating the potential of this approach for studying pathological tissue organization in three dimensions. More broadly, this framework may be valuable for studying intestinal diseases associated with altered epithelial ultrastructure and organization, including brush border remodeling in Celiac Disease and epithelial architecture and junctional defects in Inflammatory Bowel Disease.^26,27^

Although ExM currently does not achieve the resolution that some electron microscopy approaches achieve, it already occupies a useful intermediate space between conventional fluorescence microscopy and volumetric EM by combining specific antibody staining, volumetric context, and ultrastructural readout in a comparatively accessible workflow. In addition, the method supports straightforward multicolor fluorescence labeling, enabling simultaneous visualization of multiple molecular targets within intact tissue volumes. While larger expansion factors may further improve effective resolution, they also increase fluorophore dilution and expanded sample dimensions, limiting volumetric imaging depth when using high numerical aperture water immersion objectives. In this context, ∼4-fold expansion provides a favorable balance between resolution, fluorescence retention, and imaging depth, enabling imaging of intact organoids using commercially available long-working-distance water objectives with numerical apertures equal to or exceeding 1.0. Future improvements in sample preparation, anchoring chemistry, and labeling strategies may further improve structural contrast and broaden the range of ultrastructural features that can be robustly resolved.

In summary, our results establish expansion microscopy as a tool for molecularly annotated, quantitative multiscale imaging of intact intestinal epithelial systems, bridging tissue architecture, epithelial topology, and subcellular ultrastructure within a single fluorescence imaging workflow. We anticipate that this workflow will provide new insights into the organisational complexity of specialized cell types within a single imaging modality, including processes such as volumetric organelle positioning during differentiation and disease.

## ACKNOWLEDGEMENTS

This work was supported by the Netherlands Organization for Scientific Research (NWO) through the Gravitation programme IMAGINE! (Project number 24.005.009 to L.C.K and J.v.R.) and a VICI grant (VI.C.212.062 to L.C.K).

## AUTHOR CONTRIBUTIONS

T.M. established expansion methods, performed all experiments, analyzed data and wrote the manuscript; V.H. developed segmentation and analyzed data under supervision of I.S.; A.K.S. established expansion methods; D.L. and J.v.R. prepared tissue; S.J.E.S. contributed organoids and expertise; S.A.F. and K.S.V. contributed the Myo5b^fl/fl^; Vil-CreERT2 organoids; L.C.K conceived research, obtained funding, revised the manuscript and supervised the study.

## DECLARATION OF GENERATIVE AI AND AI-ASSISTED TECHNOLOGIES

During the preparation of this manuscript, the authors used ChatGPT (OpenAI) in order to correct or improve sentences. After using this tool or service, the authors reviewed and edited the content as needed and take full responsibility for the content of the publication.

## METHODS

### Organoid culture

Murine intestinal organoids carrying an mTdtomato-CAAX reporter were harvested from ROSA26-mT/mG mice.^28^ Organoids were isolated as described before.^5^ Intestinal tissue was cleaned and flushed with cold PBS. The intestine was then opened longitudinally, and the villi were removed by gentle scraping with a coverslip. After additional washing with cold PBS, the tissue was incubated in 5 mM EDTA for 30 minutes at 4 °C with gentle rolling. Crypts were subsequently isolated, passed through a 70 µm cell strainer (Greiner Bio-One), and plated. The organoid isolation was performed in accordance with the guidelines of and approved by the animal welfare committee of the Netherlands Cancer Institute.

Organoids were cultured as described in A. Krotenberg Garcia et al.^29^ Every three days, organoids were passaged by mechanically dissociating the organoid crypts using a narrowed glass pipette. The dissociated crypts were embedded in domes of Basement Membrane Extract (BME) in a ratio of 1:2 (medium : BME) on tissue culture plates that had been pre-incubated for at least 48 hours. To allow the BME to solidify, the plates were incubated upside-down for 20 minutes. Once set, the domes were submerged in growth medium containing Advanced DMEM/F-12, GlutaMAX (1x), Hepes (10 mM), Penicillin/Streptomycin (100 U/mL), N21 Supplement (1x), n-acetylcysteine (1.25 mM), human epidermal growth factor (50 ng/ml), Noggin-FC (1 nM), and R-spondin conditioned medium (10% v/v). The R-spondin1 conditioned medium was made as described in Broutier et al.^30^ Organoids were maintained in incubators at 37°C with 5% CO_2_ and 100% humidity.

The *Myo5b*^fl/fl^; Vil-CreERT2 organoids were provided by Sabine A. Fuchs and were originally made by K. Schneeberger et al.^23^ They were established from intestine-specific Cre-inducible floxed *Myo5b* mice, with *LoxP* sites placed around exon 4 of *Myo5b*. Intestine specificity was achieved by crossing *Myo5b^fl/fl^* mice to a Vil-CreERT2 line^31^, which express Cre recombinase fused to a mutated ligand-binding domain of the human estrogen receptor under control of the intestinal *Vil* promotor; Myo5b^fl/fl^; Vil-CreERT2 organoids can be induced by the addition of tamoxifen, leading to translocation of the Cre-ERT2 protein to the nucleus and recombination of the *LOXP* sites to inactivate *Myo5b*.

Organoids were passaged as described above and 1 μM of 4OH-Tamoxifen was added to the medium directly after passaging for overnight incubation. The next day the medium was replaced, and the organoids were fixed at day three after passaging.

### Organoid preparation for ExM

Round 18 mm coverslips were placed in 12-well cell culture plates. On these coverslips, 90 μL of dissociated crypts in a 1:2 (medium : BME) ratio were evenly seeded and put in the incubator at 37C with 5% CO_2_ for 20 min to solidify the BME. Afterward, the growth medium was added, and the organoids were placed in the incubator. In ∼3 days, the organoids matured and were ready for fixation.

### Fixation of organoids

The growth medium was carefully removed from the wells. The organoids were fixed using 2 ml of PHEM (60 mM Pipes, 25 mM Hepes, 10 mM EGTA, and 2 mM MgCl_2_ adjusted to pH 6.9 with NaOH) with 4% PFA for 20 to 30 min at 37 °C. This dissolved most of the BME and attached the organoids to the coverslip. To remove the fixation solution, the organoids were washed three times using 1x PBS, and either used directly for further processing or stored at 4 °C for later use.

### Pre-expansion immunostaining

The staining was adapted from the protocol described by A. Krotenberg Garcia et al.^29^ Fixed organoids were incubated in blocking buffer (1x PBS, 0.2% Triton X-100, and 5% BSA) for 30 min at room temperature. Primary antibodies were diluted in antibody buffer (1x PBS, 0.1% Triton X-100, and 2.5% BSA), and samples were incubated for 2 days at 4 °C. Samples were subsequently washed three times for 10 min in washing buffer (1x PBS, 0.1% Triton X-100). Secondary antibodies were diluted in antibody buffer and incubated with the samples overnight at 4 °C. Samples were washed three times for 10 min in washing buffer, rinsed once in 1x PBS, and either stored in 1x PBS at 4 °C or processed immediately for anchoring and expansion.

### Tissue preparation and fixation

All experiments were approved by the DEC Dutch Animal Experiments Committee (Dier Experimenten Commissie), performed in line with institutional guidelines of Dutch Cancer Institute, and conducted in agreement with Dutch law (Wet op de Dierproeven, 1996) and European regulations (Directive 2010/63/EU). After the mice were sacrificed the intestine was harvested and briefly washed in 4 °C 1xPBS without calcium. The intestine was cut into sections and put in 50 mL tubes in 4 °C PHEM buffer. The PHEM buffer was replaced by PHEM with 4% PFA and put overnight at 4 °C on a roller mixer for fixation. The PHEM was replaced by 1xPBS, and the sample was stored at 4 °C.

### Slicing Tissue

A piece (∼0.5-1 cm) of the small intestine was cut by hand from a larger fixed tissue piece from which the fat was removed. Using a paper towel excess water was removed from the tissue piece and it was embedded in a 5% agarose gel in 1xPBS. This was cut using a vibratome (Leica VT1000 S) in 200 μm thick slices. These were stored in 1xPBS at 4 °C for further processing.

### Anchoring of organoids and tissue for ExM

The 1xPBS was removed from the wells and 2 mL of anchoring solution (3% w/v Acrylamide, 1.2% w/v Formaldehyde, 1xPBS, and MQ) was added to each sample in the 12-well plate. 1xPBS was added to empty wells to prevent evaporation and a large piece of parafilm was put between the plate and the lid. Additionally, another strip of parafilm was used to seal the lid to the plate. The plate was placed in an incubator at 37 °C and 25 RPM for overnight anchoring of the sample. The following day the anchoring solution was removed, and the sample was washed with 1xPBS.

### Gelation of tissue and organoids for ExM

Samples were incubated in an inactive gelation solution (identical to the active gelation solution but lacking 4-hydroxy-TEMPO (4-HT), tetramethylethylenediamine (TEMED) and ammonium persulfate (APS)) for 1 h at 4 °C with gentle agitation to allow monomer infiltration. The inactive solution was then replaced with the corresponding active gelation solution (19% (w/v) sodium acrylate, 10% (w/v) acrylamide, 0.1% (w/v) bis-acrylamide, 0.001% (w/v) 4-HT, 0.25% (v/v) TEMED, 0.25% (w/v) APS and 1xPBS), and samples were incubated for 30 min at 4 °C with agitation. Sodium acrylate was made as described in H.Damstra et al.^32^

Coverslips (25mm diameter) were plasma-cleaned, briefly incubated with 0.1% (w/v) poly-L-lysine, and air-dried overnight before use. Tissue sections were mounted onto these poly-L-lysine-coated coverslips. These and coverslips bearing organoids were transferred to a polymerization chamber constructed from a microscope slide spacered with stacked 18 mm square coverslips (two layers for organoid samples and one layer for tissue slices) affixed with high-vacuum silicone grease to define gel thickness. The assembled chamber was incubated at 37 °C and 100% humidity for 90 min to allow polymerization. Following gelation, the polymerized gel was carefully released from the slide.

For organoid samples, the polymerized gel together with the coverslip was trimmed to fit into 2 ml tubes, whereas tissue slice gels on coverslips were transferred to glass flasks. Samples were fully immersed in pre-warmed denaturation buffer (14% (w/v) SDS, 0.5 M NaCl, 0.3 M Tris pH 8.8) to denature proteins and homogenize the embedded organoids and tissue. Organoid gels were denatured at 98 °C for 3 hours in a heating block, whereas tissue gels were denatured in an autoclave (Prestige Medical 210001) for 2 h (≈11 cycles).

Following denaturation, gels were removed from the tubes or flasks and rinsed in excess 1× PBS for 15 min with gentle agitation in 15 cm dishes. PBS was then replaced with Milli-Q water, and gels were expanded to equilibrium size through three consecutive washes with agitation. Fully expanded gels were measured to determine the expansion factor, re-equilibrated in 1× PBS to reduce size for handling and sectioned into smaller pieces for subsequent staining.

### Staining expanded samples

Expanded gels equilibrated in 1x PBS were transferred to 24-well plates and incubated in primary antibody (PA) solution (PA diluted in 1x PBS containing 0.1% (v/v) Triton X-100 and 2% (w/v) bovine serum albumin (BSA)) for 3 days at 4 °C with gentle agitation. Gels were washed three times for 15 min in 1x PBS with 0.1% (v/v) Triton X-100 at room temperature and subsequently incubated in secondary antibody (SA) solution (SA 1:500 diluted in 1x PBS, 0.1% (v/v) Triton X-100, 2% (w/v) BSA) overnight at 4 °C with agitation. After three washes (15 min each) in 1× PBS with 0.1% (v/v) Triton X-100 at room temperature, gels were incubated in tertiary antibody (TA) solution (TA 1:500 diluted in 1x PBS, 0.1% (v/v) Triton X-100, 2% (w/v) BSA) overnight at 4 °C with agitation. Gels were again washed three times for 15 min in 1x PBS with 0.1% (v/v) Triton X-100, followed by a final rinse in 1× PBS. For total protein labeling, gels were incubated with NHS-ester dye (with or without DAPI) in 1x PBS for ≥2 h at room temperature, protected from light. Samples were washed three times in 1x PBS and either imaged immediately or stored in 1x PBS at 4 °C until imaging. Antibodies and dyes, including working concentrations, are listed in the Key Resources Table.

### Confocal Imaging of Expanded Samples

ExM samples were cut to size and attached on poly-L-lysine coated coverslips No. 1.5, which were mounted in Attofluor cell chambers (Thermo Fisher Scientific A7816) and submerged in Milli-Q. The expanded organoids were imaged using a Zeiss LSM980 Airyscan-2 with auto-immersion and a 40x water-immersion objective with a numerical aperture of 1.1 and a working distance of 0.62 mm (421867-9970-000). The laser lines used were 405/488/639 nm. Given the larger size of expanded samples, using a long-working-distance objective is key for imaging whole expanded organoids. Images were acquired in SR Airyscan mode. Raw images were processed using Zeiss 2D/3D Airyscan processing.

### Image segmentation and quantification

Quantitative analysis of the total protein stain (NHS-Atto643), the membrane marker (CAAX-tdTomato), and the nucleus (DAPI) was performed on a three-dimensional image volume containing a complete intestinal crypt section, corresponding to 1051x1051x1051 voxels (336.3x345.8x345.8µm). Pre-processing was done within Zeiss Zen 3.12 using the option Stack-correction followed by Airyscan processing (with the parameter “super-resolution=5.2, 3D”).

The image was manually thresholded to generate a binary mask of the crypt. The largest connected component was extracted, and morphological hole-filling was applied to obtain include missing villus and Paneth cell regions.

The villus region was extracted with SAM2^33^ using 5 bounding box prompts and the provided pre-trained model “sam2.1_hiera_l” (with the Z-axis as the required temporal dimension). Manual refinements of the apical Paneth cell membrane, as well as manual improvements mentioned below, were done using the layer control in the Napari GUI.

Nuclei were segmented using Cellpose^34^ with the pre-trained model “CellposeSAM” and the following model parameters: “diameter: 120”, “do3D True”, “flow_threshold 0.4”, “cell probability threshold -2.0” and “flow smoothing 3D sigma 2.0”. The chromosomes of dividing cells were manually annotated based on DAPI signal intensity.

CAAX-tdTomato was further processed with two iterations of directional filtering^35^ (taking the maximum value over median values along 129 oriented line segments with length of 17 voxels) using a custom 3D Cupy^36^ implementation, followed by rolling ball background subtraction^37^ with the radius set to 50 voxels.

The nuclei and crypt segmentations together with the processed CAAX-tdTomato signal were used to create automated cell segmentation using the seeded watershed, implemented using MCP_flexible module from Sci-kit image.^37^ The seeds were defined by the nuclei centers of mass, and the cost function was defined as: zero inside the nucleus, CAAX-tdTomato signal intensity to the power of 5 inside the crypt and a maximum penalty outside the crypt.

The segmentation was then transformed into a cylindrical coordinate system to aid manual data curation and improve the accuracy of the cell segmentation. As such, apical-basal cross-sections of all cells become clearly visible and aligned within one image plane. The curated planes were interpolated using a Napari label interpolator (https://github.com/brisvag/napari-label-interpolator), particularly to improve accuracy near the apical membrane. Any remaining unlabeled voxels within the crypt were assigned to the closest cell. These segmentation masks were then transformed back into the original Cartesian coordinate system. For each cell, all the quantitative measurements were computed jointly (in parallel) using Dask-image labeled_comprehension^38^ to minimize memory (re-)distribution overhead, and speedup the computation.

This framework enables custom metrics to be explored, particularly to capture the characteristic elongated shape of ISCs. This included the principal axes derived metrics, as well as metrics from elliptical^39^ and rectangular^40^ shape approximations.

## KEY RESOURCES TABLE

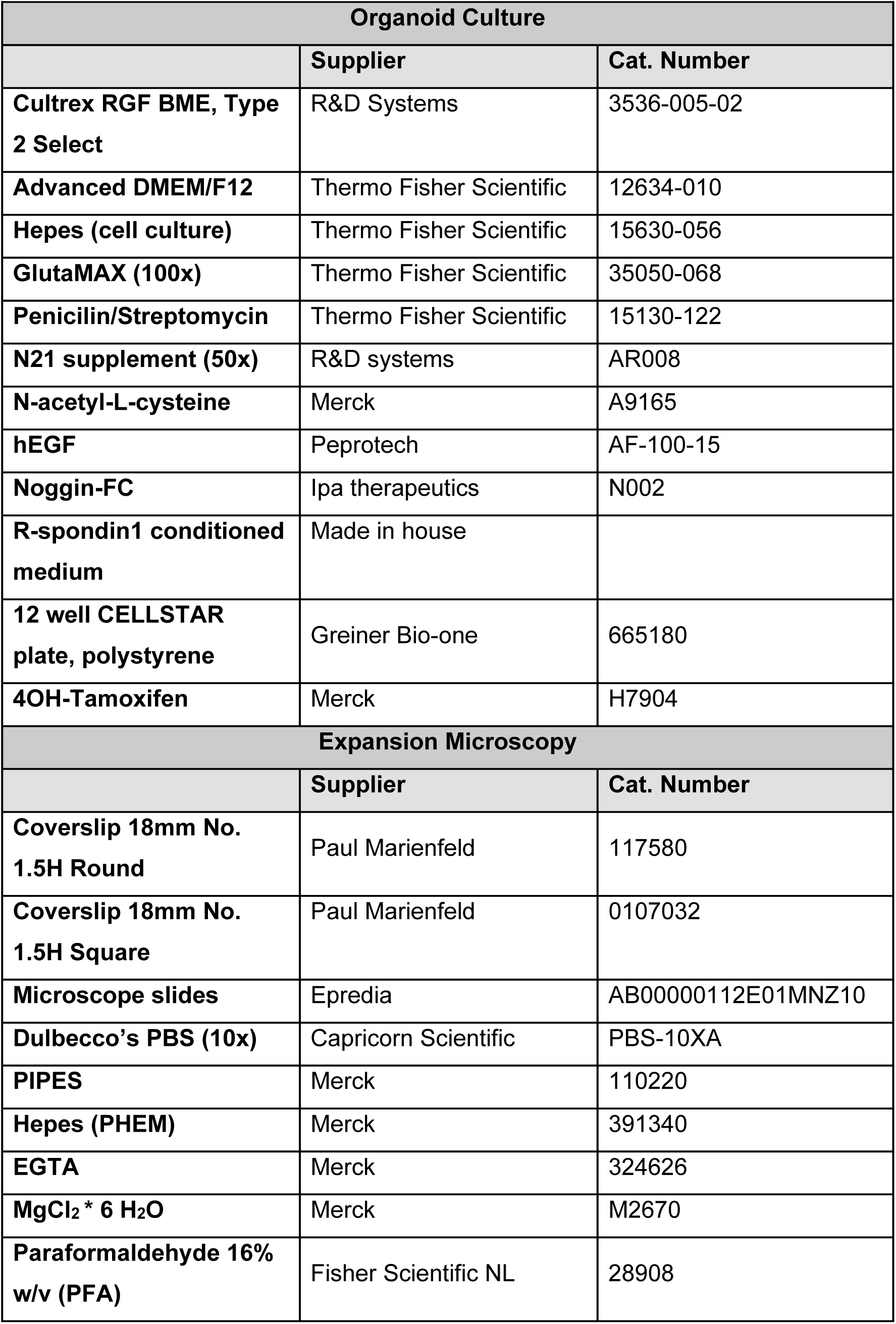
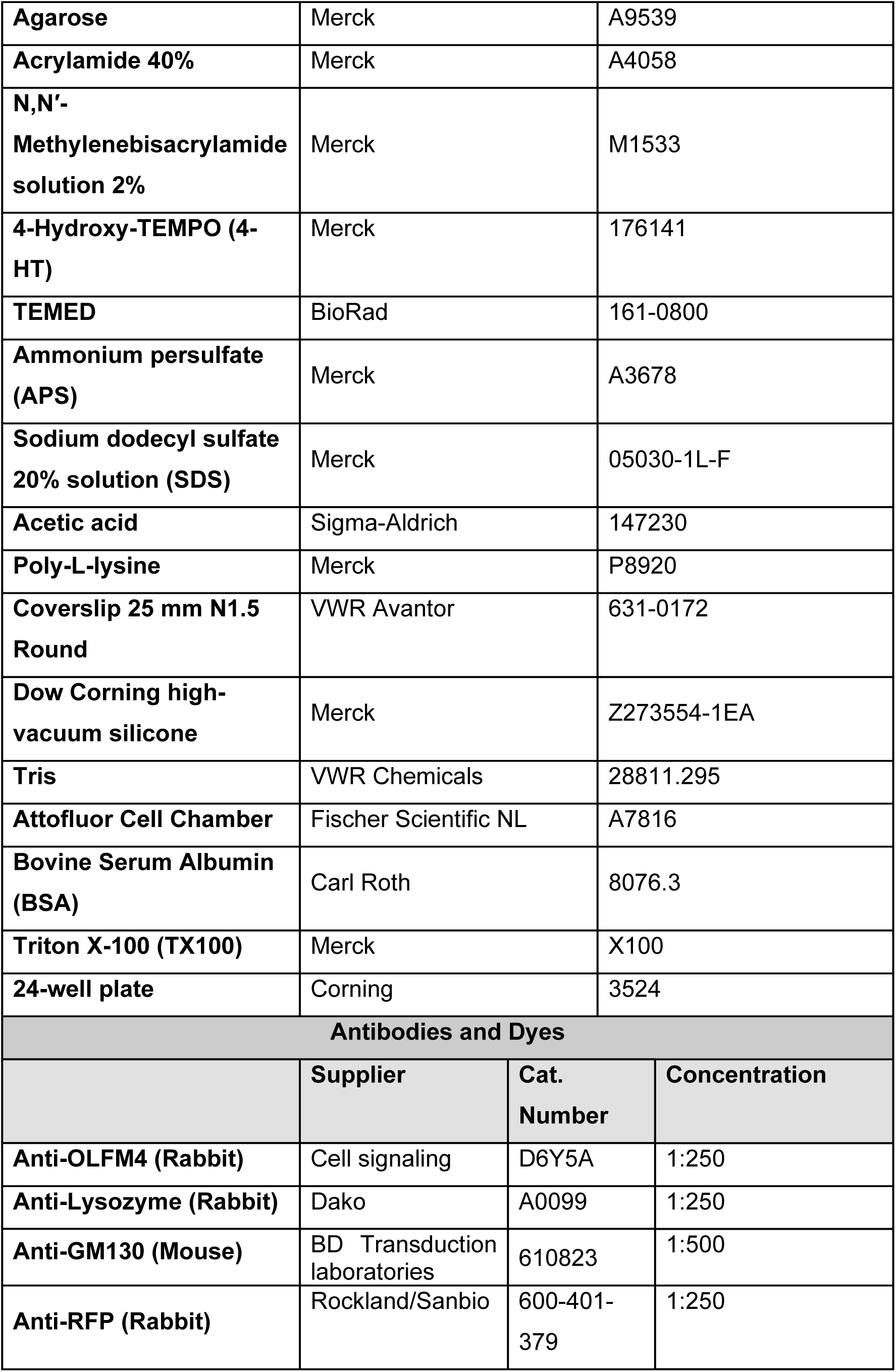
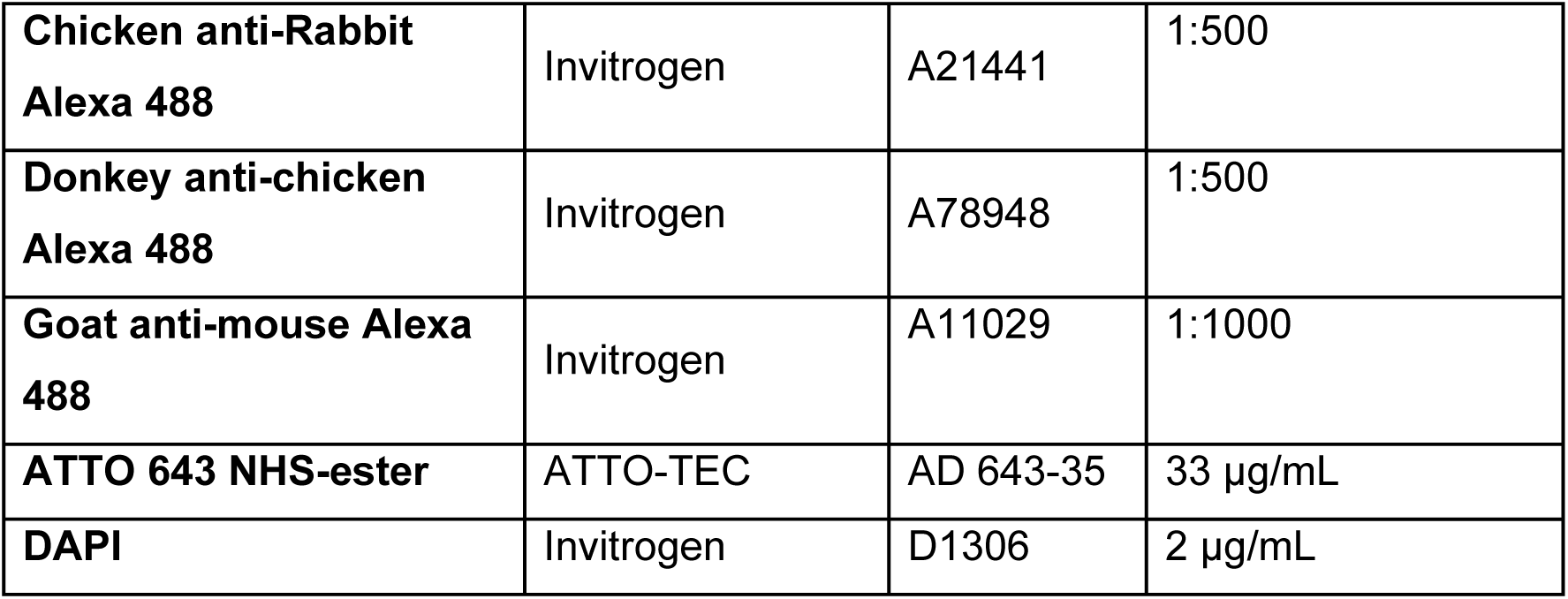

| **Organoid Culture** |  |  |
| --- | --- | --- |
|  | **Supplier** | **Cat. Number** |
| **Cultrex RGF BME, Type 2 Select** | R&D Systems | 3536-005-02 |
| **Advanced DMEM/F12** | Thermo Fisher Scientific | 12634-010 |
| **Hepes (cell culture)** | Thermo Fisher Scientific | 15630-056 |
| **GlutaMAX (100x)** | Thermo Fisher Scientific | 35050-068 |
| **Penicilin/Streptomycin** | Thermo Fisher Scientific | 15130-122 |
| **N21 supplement (50x)** | R&D systems | AR008 |
| **N-acetyl-L-cysteine** | Merck | A9165 |
| **hEGF** | Peprtech | AF-100-15 |
| **Noggin-FC** | Ipa therapeutics | N002 |
| **R-spondin1 conditioned medium** | Made in house |  |
| **12 well CELLSTAR plate, polystyrene** | Greiner Bio-one | 665180 |
| **4OH-Tamoxifen** | Merck | H7904 |
| **Expansion Microscopy** |  |  |
|  | **Supplier** | **Cat. Number** |
| **Coverslip 18mm No. 1.5H Round** | Paul Marienfeld | 117580 |
| **Coverslip 18mm No. 1.5H Square** | Paul Marienfeld | 0107032 |
| **Microscope slides** | Epredia | AB00000112E01MNZ10 |
| **Dulbecco's PBS (10x)** | Capricorn Scientific | PBS-10XA |
| **PIPES** | Merck | 110220 |
| **Hepes (PHEM)** | Merck | 391340 |
| **EGTA** | Merck | 324626 |
| **MgCl<sub>2</sub> \* 6 H<sub>2</sub>O** | Merck | M2670 |
| **Paraformaldehyde 16% w/v (PFA)** | Fisher Scientific NL | 28908 |

|  |  |  |  |
| --- | --- | --- | --- |
| Agarose | Merck | A9539 |  |
| Acrylamide 40% | Merck | A4058 |  |
| N,N'-Methylenebisacrylamide solution 2% | Merck | M1533 |  |
| 4-Hydroxy-TEMPO (4-HT) | Merck | 176141 |  |
| TEMED | BioRad | 161-0800 |  |
| Ammonium persulfate (APS) | Merck | A3678 |  |
| Sodium dodecyl sulfate 20% solution (SDS) | Merck | 05030-1L-F |  |
| Acetic acid | Sigma-Aldrich | 147230 |  |
| Poly-L-lysine | Merck | P8920 |  |
| Coverslip 25 mm N1.5 Round | VWR Avantor | 631-0172 |  |
| Dow Corning high-vacuum silicone | Merck | Z273554-1EA |  |
| Tris | VWR Chemicals | 28811.295 |  |
| Attofluor Cell Chamber | Fischer Scientific NL | A7816 |  |
| Bovine Serum Albumin (BSA) | Carl Roth | 8076.3 |  |
| Triton X-100 (TX100) | Merck | X100 |  |
| 24-well plate | Corning | 3524 |  |
| Antibodies and Dyes |  |  |  |
|  | Supplier | Cat. Number | Concentration |
| Anti-OLFM4 (Rabbit) | Cell signaling | D6Y5A | 1:250 |
| Anti-Lysozyme (Rabbit) | Dako | A0099 | 1:250 |
| Anti-GM130 (Mouse) | BD Transduction laboratories | 610823 | 1:500 |
| Anti-RFP (Rabbit) | Rockland/Sanbio | 600-401-379 | 1:250 |
| **Chicken anti-Rabbit**<br>**Alexa 488** | Invitrogen | A21441 | 1:500 |
| **Donkey anti-chicken**<br>**Alexa 488** | Invitrogen | A78948 | 1:500 |
| **Goat anti-mouse Alexa**<br>**488** | Invitrogen | A11029 | 1:1000 |
| **ATTO 643 NHS-ester** | ATTO-TEC | AD 643-35 | 33 µg/mL |
| **DAPI** | Invitrogen | D1306 | 2 µg/mL |

## SUPPLEMENTARY FIGURES

**Supplementary Figure 1.**
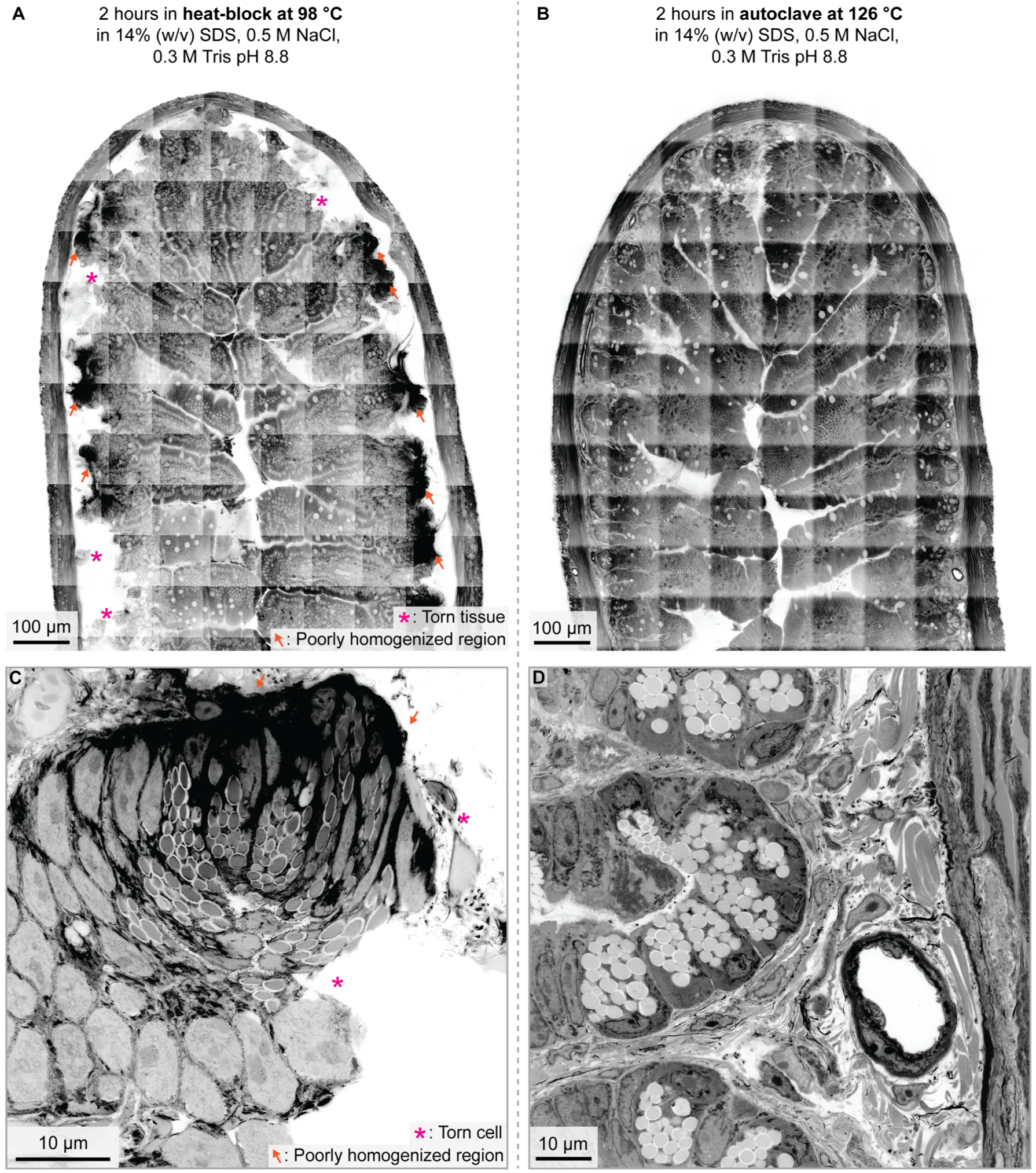
Optimization of homogenization conditions for intestinal tissue sections. **A)** Tissue integrity is compromised following heat-block homogenization. Torn regions are indicated by magenta asterisks, while poorly homogenized protein-dense regions are indicated by orange arrows. **B)** Tissue architecture and continuity are preserved following autoclave homogenization. **C)** Higher-magnification view of an intestinal crypt after heat-block homogenization showing disrupted cellular integrity. A torn cell is indicated by a magenta asterisk and a poorly homogenized protein-dense region by an orange arrow. **D)** Higher-magnification view of an intestinal crypt after autoclave homogenization demonstrating preservation of cellular and tissue integrity within and surrounding the crypt. Scale bars are corrected for the expansion factor. All images show NHS-ester Atto 643 total protein staining.

**Supplementary figure 2.**
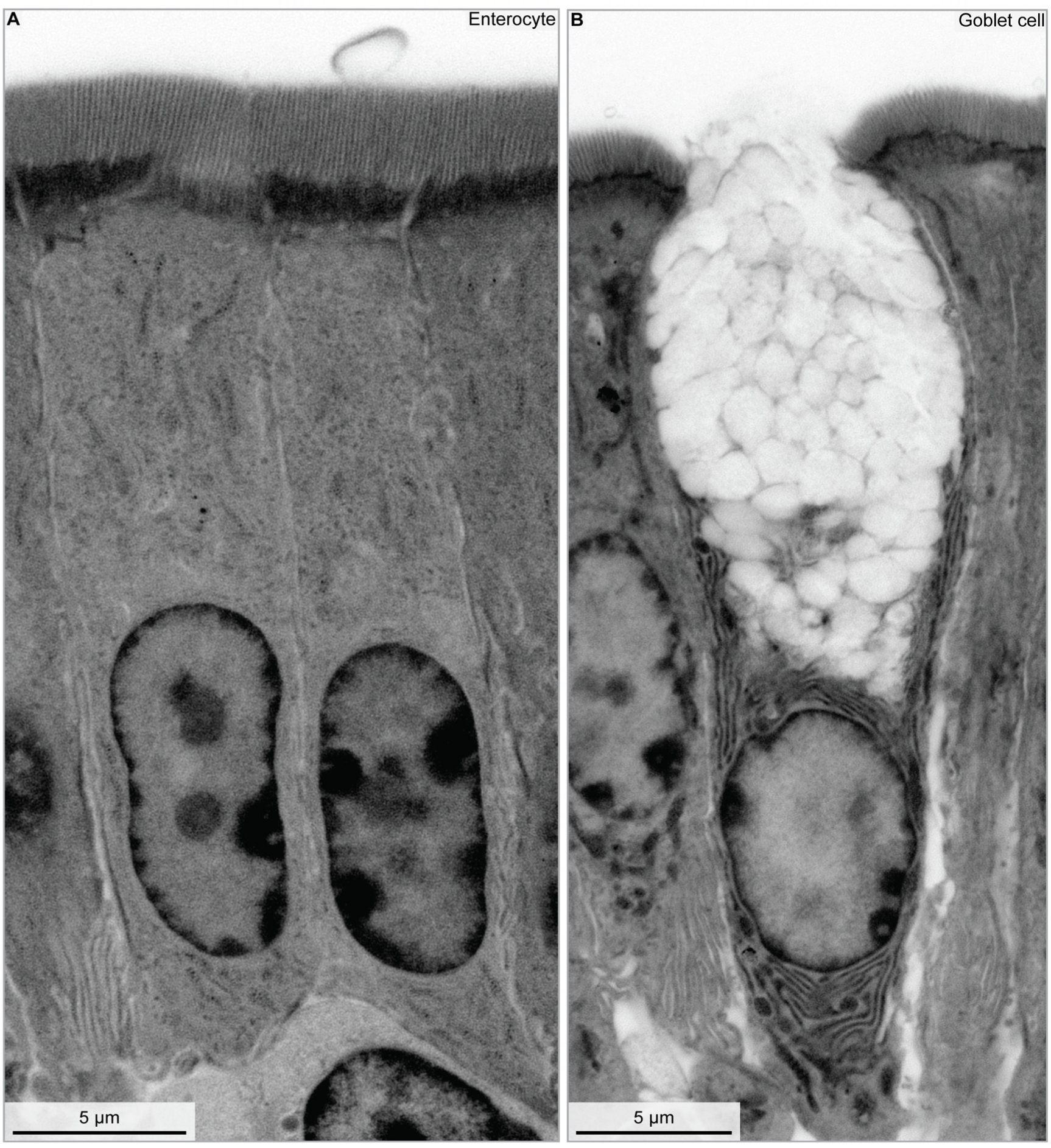
Enlarged views of subcellular contrast in enterocyte and goblet cell. **A)** Enlarged view of the enterocyte shown in Fig. 1, highlighting subcellular vesicles, lateral membrane interdigitations, and a densely packed brush border. **B)** Enlarged view of the goblet cell shown in Fig. 1, highlighting protein-dense subcellular organelles. Scale bars are corrected for the expansion factor. All images show NHS-ester Atto 643 total protein staining.

**Supplementary figure 3.**
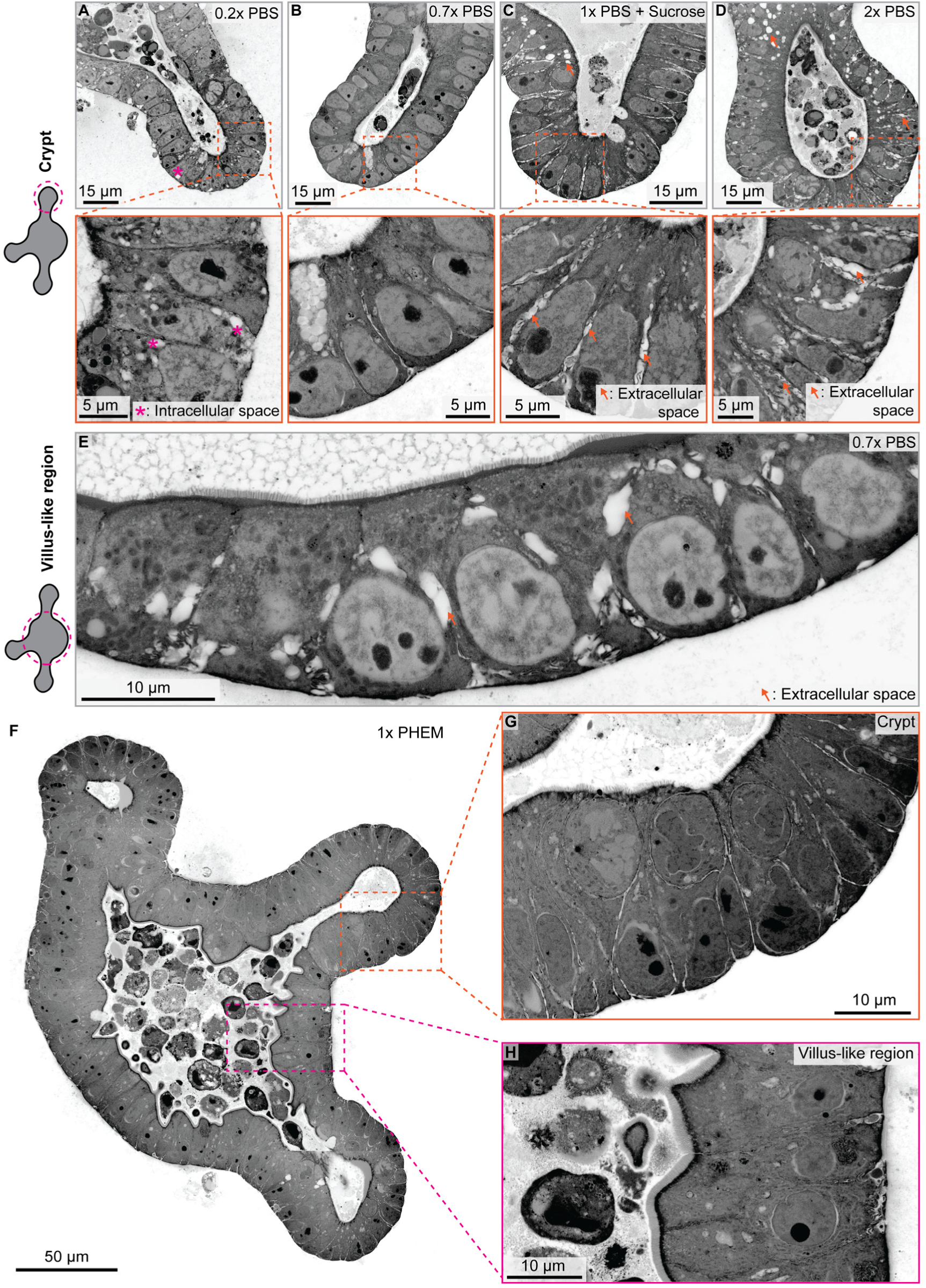
Impact of fixation buffer composition on epithelial integrity. **A–D)** Representative images of organoids fixed with 4% PFA prepared in 0.2× PBS, 0.7× PBS, 1× PBS supplemented with sucrose, and 2× PBS. Asterisks indicate intracellular spaces and arrows indicate extracellular spaces. Fixation in 0.7× PBS reduced the number of intracellular and intercellular swellings within the crypt epithelium. **E)** Villus-like region of an organoid fixed with 4% PFA in 0.7× PBS, showing the presence of intercellular swelling within the epithelium. **F)** Optical section of a whole organoid fixed with 4% PFA in PHEM buffer. **G)** Enlarged view of the crypt region shown in (**F**), revealing an intact epithelial layer with limited extracellular spaces between cells. **H)** Enlarged view of the villus-like region shown in (**F**), revealing an intact epithelial layer with limited extracellular spaces between cells. Scale bars are corrected for the expansion factor. All images show the NHS-ester Atto-643 total protein stain.

## SUPPLEMENTARY VIDEOS

**Supplementary Video 1 | Multiscale visualization of an expanded small intestine tissue slice.**

Overview of an expanded mouse small intestine tissue section spanning the villus tip to the crypt. The video progressively zooms from the tissue scale to individual enterocytes at the villus tip to visualize brush border microvilli, before navigating across the epithelium toward the crypt. At the crypt, the image stack is rendered in three dimensions to illustrate the volumetric organization of the tissue. Corresponds to Fig. 1B-E.

**Supplementary Video 2 | Three-dimensional segmentation of the intestinal organoid crypt.**

A z-stack through an expanded intestinal organoid crypt illustrating the segmentation workflow. The epithelial layer is first segmented, followed by segmentation of individual nuclei based on DAPI staining and individual cell bodies based on membrane-localized CAAX-tdTomato fluorescence. Corresponds to Fig. 4A.

**Supplementary Video 3 | Three-dimensional rendering of a segmented intestinal organoid crypt.**

Three-dimensional rendering of a segmented intestinal organoid crypt. The rendering is sequentially sectioned to visualize the quality of the segmentation, the continuity of the epithelial layer, and the three-dimensional organization of individual segmented cells. Corresponds to Fig. 4C

**Supplementary Video 4 | Three-dimensional visualization of thin elongated cells within an intestinal organoid crypt.**

Three-dimensional rendering of the thin elongated cells highlighted in Fig. 4D. Cell bodies are shown in orange, interfaces with Paneth cells in cyan, basal surfaces in green, and apical surfaces in magenta. The video concludes with a representative thin elongated cell (orange) wrapping around a Paneth cell (magenta), with their shared interface highlighted in cyan. Corresponds to Fig. 4D-G.

**Supplementary Video 5 | Three-dimensional visualization of a scutoid-like cell within an intestinal organoid crypt.**

Three-dimensional rendering of four neighboring epithelial cells forming a scutoid-like cell configuration. The video illustrates the change in cell neighbors along the apico-basal axis. Three of the cells are subsequently removed to reveal the characteristic geometry of the remaining scutoid-like cell. Corresponds to Fig. 4I-L.

**Supplementary Video 6 | Three-dimensional visualization of a microvillus inclusion.**

Three-dimensional rendering of a microvillus inclusion within an epithelial cell. The rendering highlights the microvilli contained within the inclusion, the reduced apical brush border, and the accumulation of subapical vesicles. Corresponds to Fig. 5C.

