## Supplementary figures and images for "Multiscale three-dimensional ultrastructural mapping of intestinal tissues and organoids"

### Figure S1

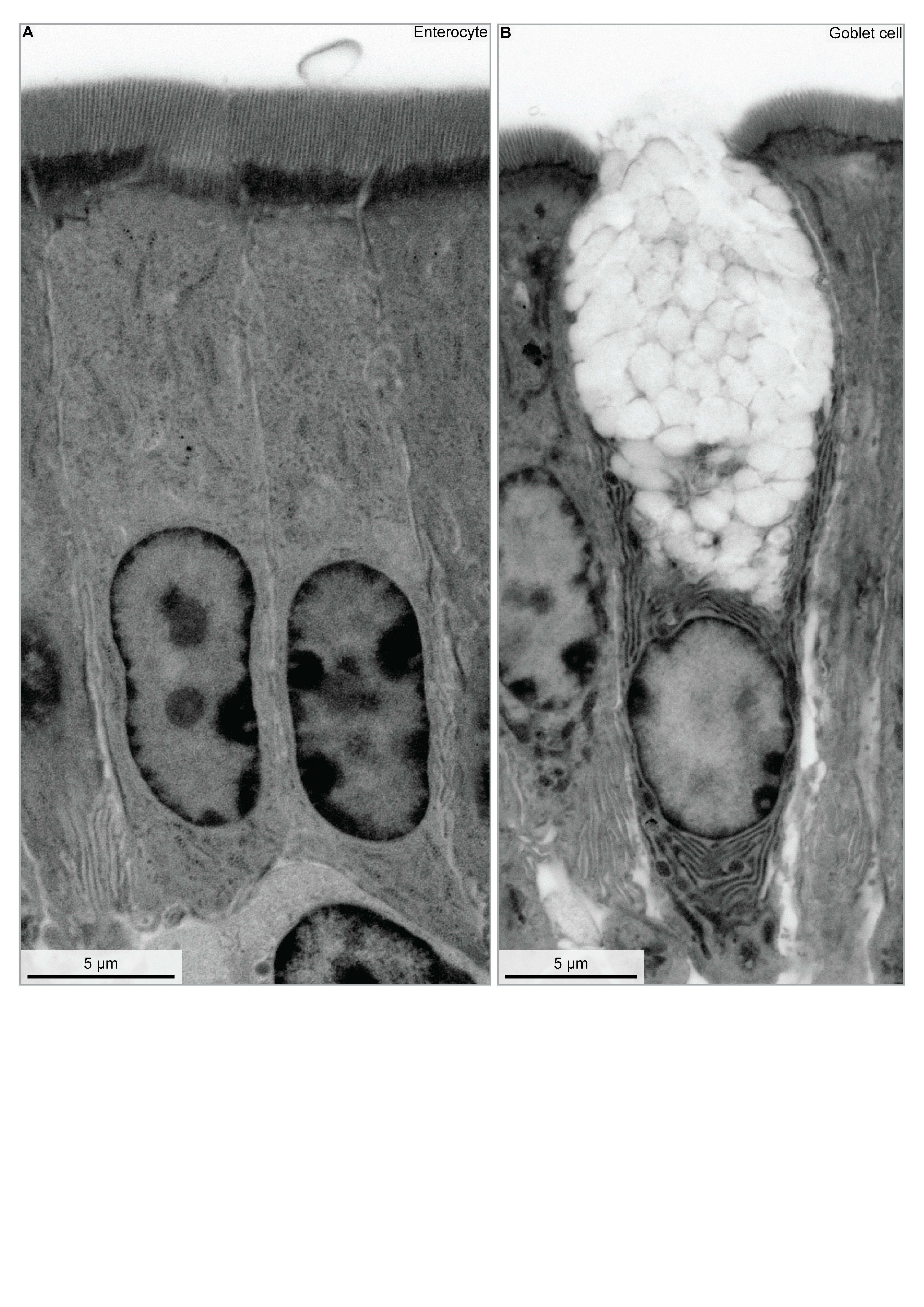

### Figure S2

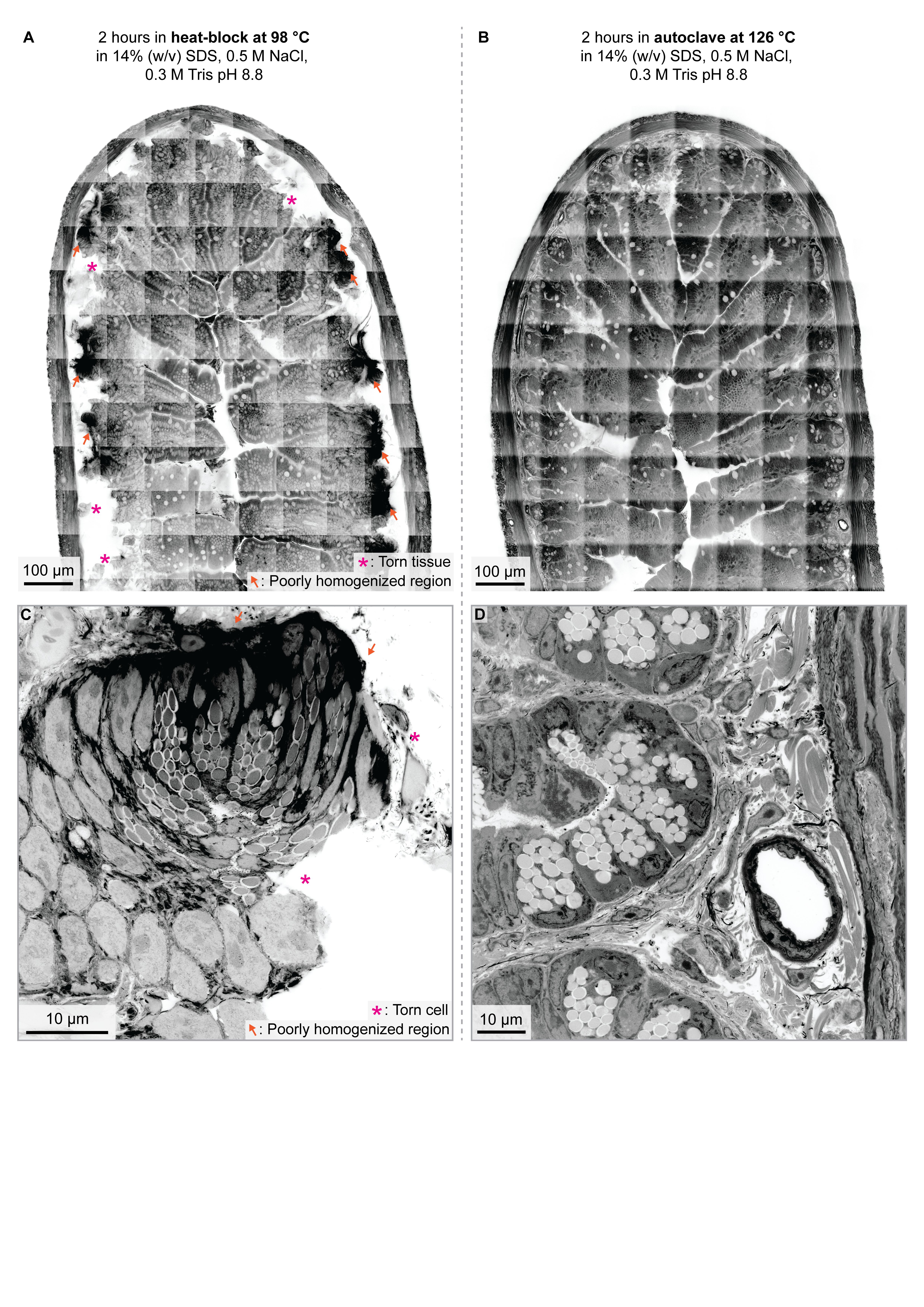

### Figure S3

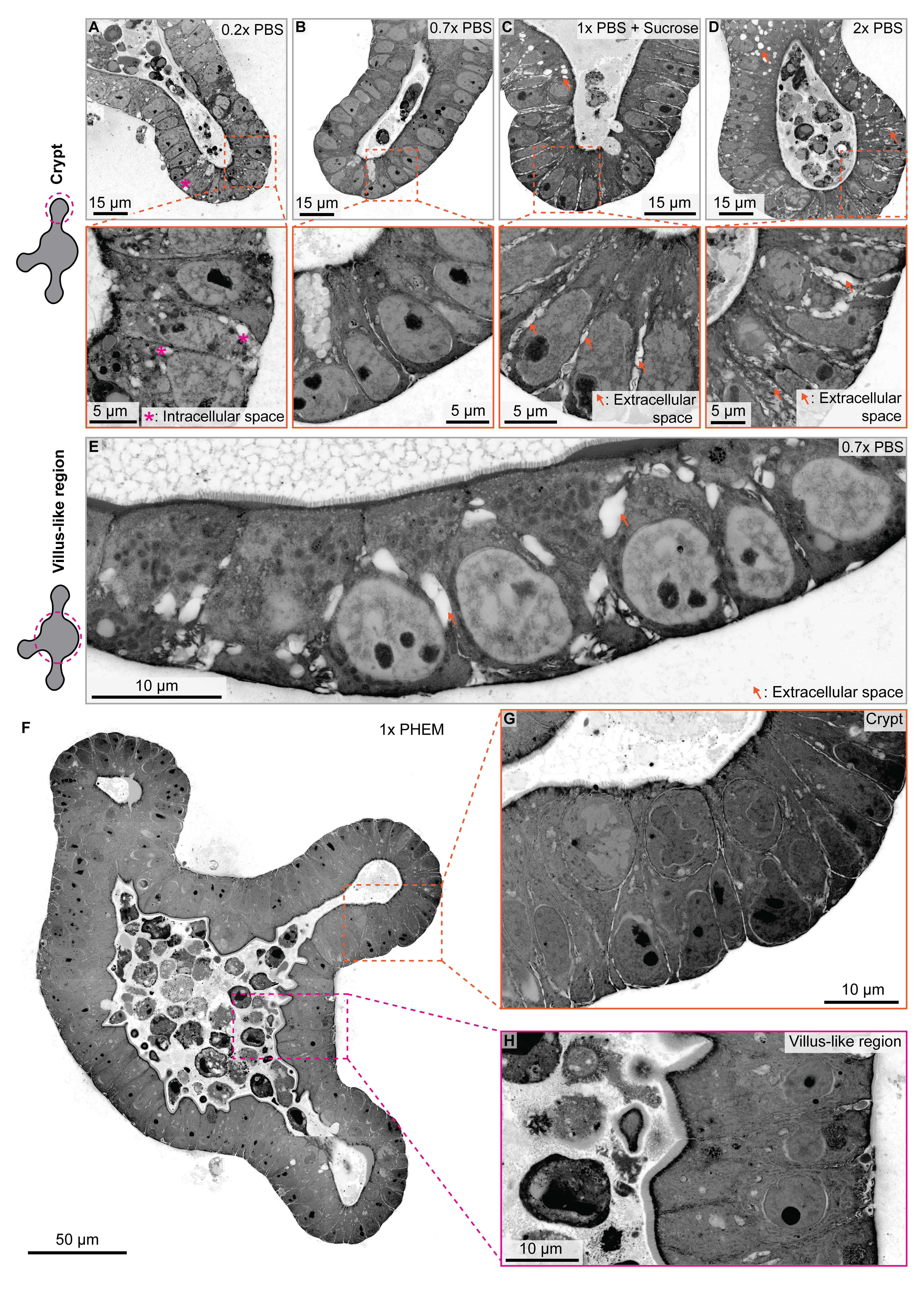
